# Cryo-EM Structure of the 50S-HflX Complex Reveals a Novel Mechanism of Antibiotic Resistance in *E. coli*

**DOI:** 10.1101/2022.11.25.517942

**Authors:** Damu Wu, Yuhao Dai, Ning Gao

## Abstract

Bacterial HflX is a conserved ribosome-binding GTPase involved in splitting ribosomal complexes accumulated under stress condition. However, the atomic details of its ribosomal interaction remain to be elucidated. In this work, we present a high-resolution structure of the *E. coli* 50S subunit bound with HflX. The structure reveals highly specific contacts between HflX and the ribosomal RNA, and in particular, an insertion loop of the N-terminal domain of HflX is situated in the peptidyl transferase center (PTC) and makes direct interactions with PTC residues. Interestingly, this loop displays steric clash with a few PTC-targeting antibiotics on the 50S subunit, such as chloramphenicol. Deletion of *hflX* results in hypersensitivity to chloramphenicol treatment, and a loop residue G154 of HflX is important for the observed chloramphenicol resistance. Overall, our results suggest that HflX could be a general stress response factor that functions in both stalled ribosome splitting and PTC antibiotic displacing.

## Introduction

HflX is a universally conserved GTPase in all domains of life (Verstraeten et al., 2011), with its eukaryotic homologues present in organelles, such as mitochondria (Gianfrancesco et al., 1998). Early studies showed that HflX was able to associate with the 50S ribosomal subunit (Blombach et al., 2011; Fischer et al., 2012; Jain et al., 2009; Polkinghorne et al., 2008), and its low intrinsic GTPase activity could be greatly stimulated by the 50S subunit (Jain et al., 2009; Shields et al., 2009) or the programmed 70S ribosome (Shields et al., 2009), implicating a possible role in ribosome-related functions. The *hflX* gene in *E. coli* is localized to the *hflA* locus (Banuett and Herskowitz, 1987; Noble et al., 1993), an operon that is under control of a heat sensitive promoter. The mRNA level of *hflX* was shown to increase upon heat shock (Carruthers and Minion, 2009; Chuang and Blattner, 1993; Richmond et al., 1999; Zhang et al., 2015), and the knockout strain of Δ*hflX* exhibited a severe growth defect under high temperature (Zhang et al., 2015). *E. coli* HflX was later found to possess an intrinsic ribosome-splitting activity, capable of disassembling vacant as well as certain mRNA-containing 70S ribosomes into subunits (Coatham et al., 2016; Zhang et al., 2015). The heat shock is known to induce pausing of translating ribosomes at early elongation steps in both eukaryotes and prokaryotes (Goenka et al., 2018; Shalgi et al., 2013; Zhang et al., 2017), which frequently resulted in the formation of non-productive, stalled ribosomes through various mechanisms (Doma and Parker, 2006; Hayes and Sauer, 2003). Therefore, HflX was proposed to be responsible for splitting and recycling stalled non-productive 70S ribosomal complexes accumulated under heat-shock conditions (Zhang et al., 2015). The endogenous substrates of HflX is not limited to heat-induced 70S ribosomal complexes, but also includes hibernating 100S ribosomes (Basu and Yap, 2017) and macrolide stalled ribosomes (Rudra et al., 2020).

Previously, we have reported a medium-resolution cryo-EM structure of the 50S subunit bound with HflX and revealed the molecular mechanism of its ribosome-splitting activity (Zhang et al., 2015). *E. coli* HflX is composed of three domains, a relatively basic N-terminal domain (NTD), a central GTPase domain (GD) and a small C-terminal domain (CTD). In the structure, the NTD of HflX interacts with the 23S rRNA through its basic residues, while the CTD interacts with a ribosomal protein uL11. Compared with the free 50S subunit, the association of HflX induces a large structural displacement of the Helix 69 of the 23S rRNA (H69), in a conformation incompatible with the binding of the 30S subunit. Very interestingly, the structure also showed that the NTD extends to the peptidyl-transferase center (PTC) in a strikingly similar way as release factors do, and a long loop in the NTD (N-loop) may have interactions with the PTC rRNA residues (Zhang et al., 2015).

More recently, it has been reported that the homolog of HflX mediates the antibiotic resistance of *L. monocytogenes* to lincosamides and of *M. abscessus* to macrolides/ lincosamides (Duval et al., 2018; Rudra et al., 2020). Based on the structure of the *E. coli* 50S-HflX complex (Zhang et al., 2015), the PTC-interacting N-loop of HflX is near or directly overlaps with a few PTC-target antibiotics on the 50S subunit (Wilson et al., 2020). Thus, HflX could has another role in translation regulation by modulating the actions of PTC-targeting antibiotics. More interestingly, this loop is in the most diverse region of HflX, varying in both the length and amino acid composition, suggesting that observed HflX-mediated resistance in pathogenic bacteria could be species-specific and antibiotic-specific (Wilson et al., 2020). The resistance to macrolide-lincosamide antibiotics in mycobacteria appeared to be coupled to the splitting activity of HflX (Rudra et al., 2020). However, whether HflX has a general role in mediating antibiotic resistance was not determined. The PTC-interacting N-loop, which could potentially regulate the binding or directly dislodge antibiotics from the PTC, was not resolved in the previous structure, and the atomic details between HflX and the critical PTC residues remain to be elucidated.

In the present work, we present a high-resolution (2.5 Å) cryo-EM structure of the *E. coli* 50S-HflX complex in the presence of GMPPNP. The structure reveals specific interactions between the N-loop of HflX and a few functionally important residues of the PTC. The R153 and G154 of the loop shows steric clash with chloramphenicol (CHL) on the 50S subunit. Consistently, the knockout of *hflX* gene or mutations of N-loop residues resulted in increased sensitivity to the CHL treatment. These data suggest that HflX has a general function in antibiotic resistance, and the species-specific N-loop might account for the observed differences of antibiotic resistance in different species.

## Results

### Overall structure of the 50S-HflX complex at 2.5-Å resolution

To understand interaction details of HflX with the PTC, we assembled the 50S-HflX complex in vitro in the presence of excessive GMPPNP, and obtained a cryo-EM density map at a final resolution 2.5 Å (Fig. 1a and Supplementary Fig. 1). The high-resolution map enabled atomic modelling of all the three domains of HflX, including the previously uncharacterized N-loop (Fig. 1b-c and Supplementary Fig. 2). The GMPPNP is also clearly seen in the nucleotide binding pocket of the GD (Supplementary Fig. 2a).

**Fig. 1.**
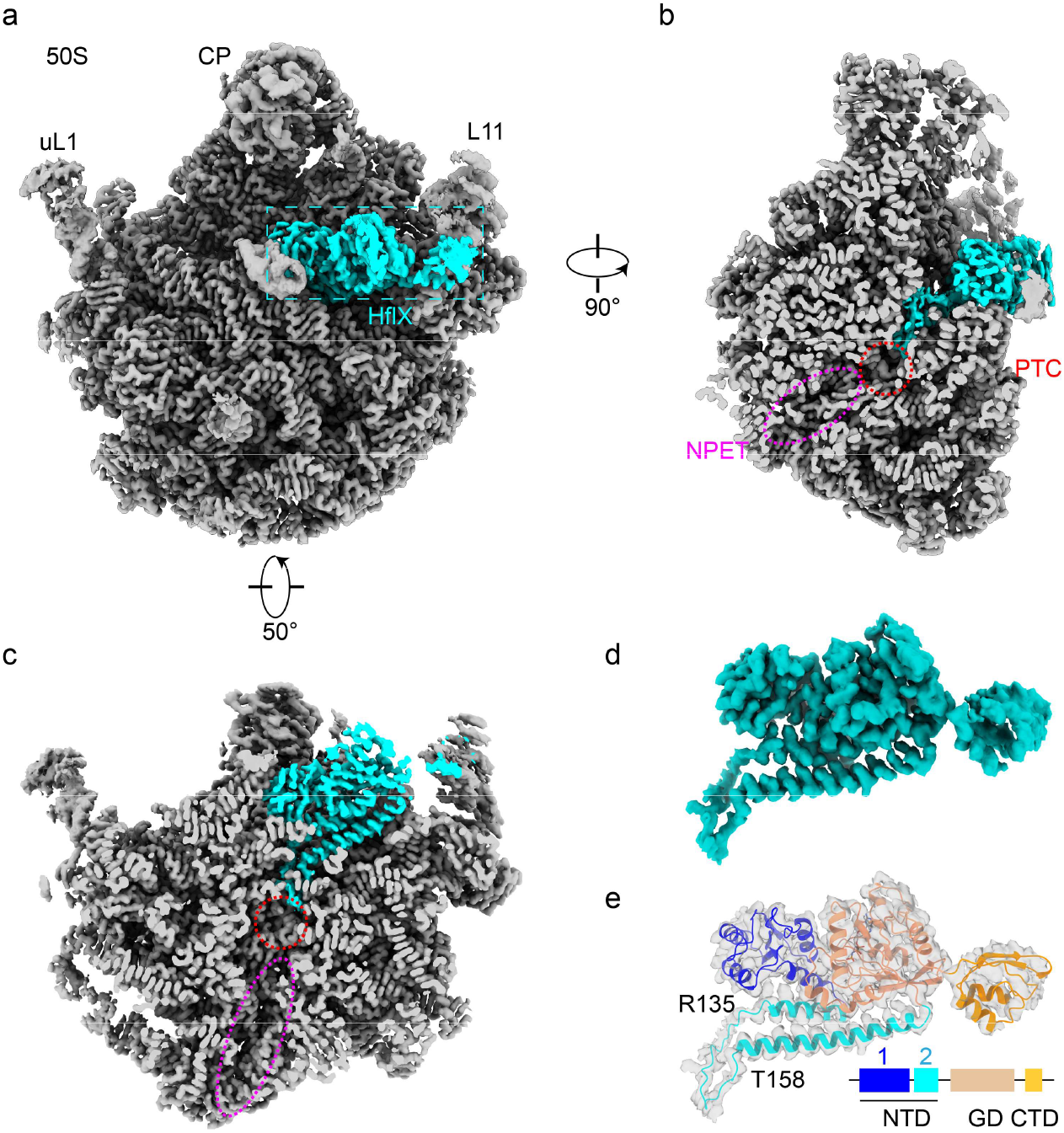
Cryo-EM structure of the 50S-HflX complex. **a**, Overall cryo-EM density map of the 50S subunit (gray) in complex with HflX (cyan). **b-c**, Sectional view of the HflX-50S complex. The regions indicated by a red circle and a magenta oval represents the peptidyl transferase center (PTC) and the nascent polypeptide exit tunnel (NPET). **d-e**, The segmented density map (d) of HflX, superimposed with the atomic model (e).

Consistent with our previous data (Zhang et al., 2015), the NTD of HflX has extensive interactions with rRNA helices. Of the two subdomains, subdomain 1 (1-115) is a globular form with four α-helices and three β-strands, while the subdomain 2 (116-197) consists of two long helices connected by a long loop (Fig. 1d-e). This N-loop of HflX (135-158) exactly protrudes towards the PTC and makes contacts with the rRNA residues at the entry of the nascent peptide tunnel (Fig. 1b-c).

### Extensive interactions of the NTD with rRNA helices at the PTC

In the cryo-EM structure, the NTD of HflX interacts extensively with the rRNA helices around the PTC (Fig. 2a). Both the subdomain 1 and subdomain 2 participate the direct interaction with the phosphate backbone of the rRNA helices. Many basic residues ofsubdomain1, such as R49, K50, K55, K62, R90 and R94, form a positively charged patch and interact with the phosphate backbone of H69 and H71 (Supplementary Fig. 2c). The second α-helix of subdomain 2, roughly in parallel with H89, forms successive polar interactions with the backbone of H89, through its basic residues, such as R164, R165, R168, R170, R177, R180, K183, R185 and R192 (Supplementary Fig. 2d).

**Fig. 2.**
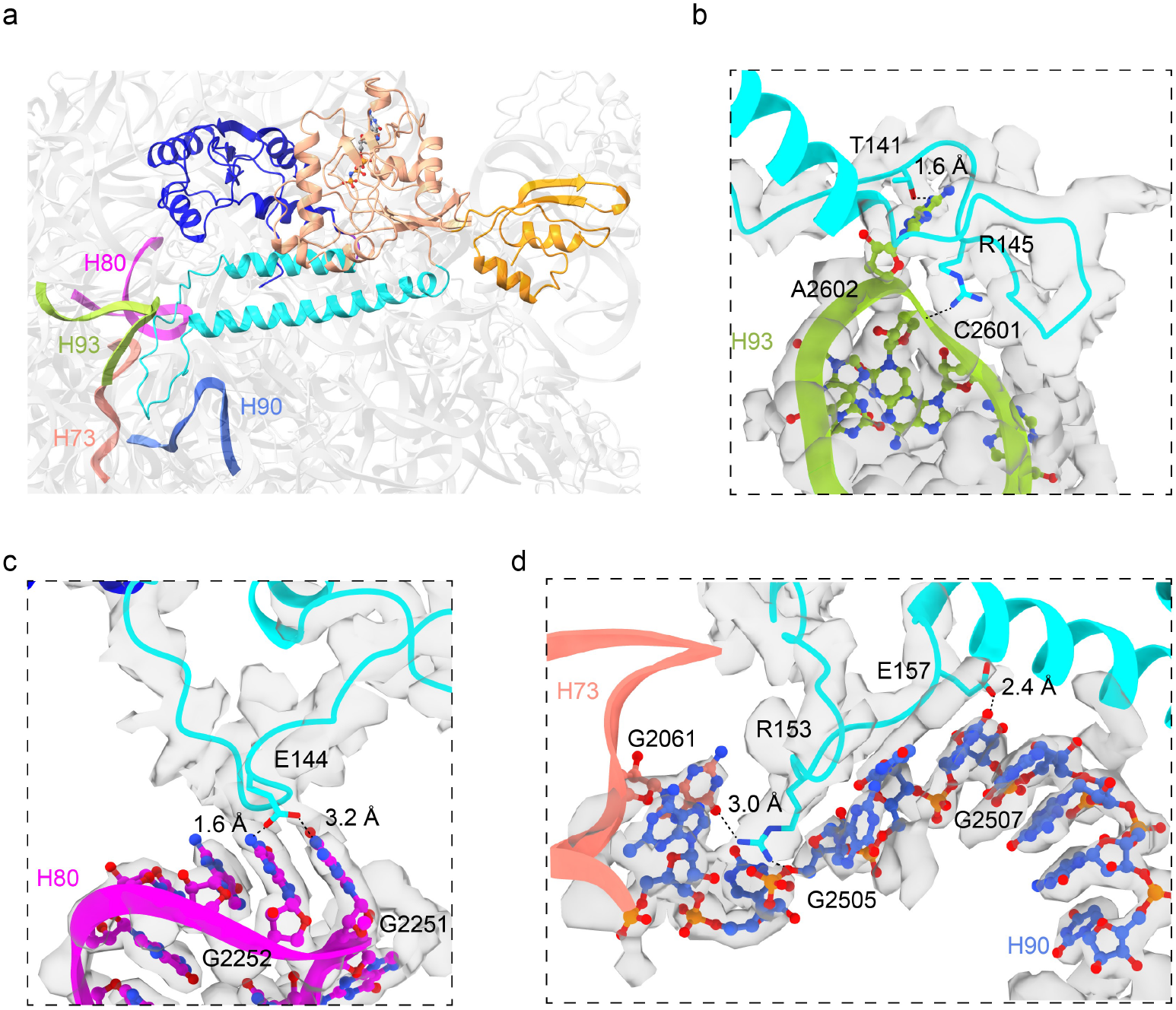
The N-loop of HflX interacts extensively with the rRNA helices around the PTC. **a**, The atomic model of the HflX-50S complex. H73, H80, H90 and H93 are colored salmon, magenta, royal blue and green, respectively. **b-d**, Zoom-in views of the interactions between the N-loop of HflX and H93, H80, H73 and H90, respectively. Both the atomic model and cryo-EM density map are shown and colored as indicated.

Very importantly, the N-loop (135-158) extends to the PTC and has strong interactions with rRNA residues of PTC helices (Fig. 2a). Firstly, on the rRNA H93, T141 interacts with the flipped base of A2602 via hydrogen bond, and R145 interacts with the phosphate backbone of G2601 through electrostatic interaction (Fig. 2b). A2602 is a universally conserved residue essential for the peptide release from the P-site tRNA (Polacek et al., 2003). Secondly, E144 of HflX interacts with the bases of G2551 and G2552 from the ribosomal A-loop (H80) via hydrogen bonds (Fig. 2c). G2553 is also universally conserved, which base-pairs to C75 of the A-site tRNA (Kim and Green, 1999) and is critically required for the A-site tRNA accommodation and following trans-peptidyl reaction (Polacek and Mankin, 2005). Thirdly, R153 points to G2505 of H90, and interacts with the phosphate backbone of G2505 and hydrogen bonds with the base of G2061 of H73 (Fig. 2d). In addition, E157 is seen to interact with G2507. Notably, G2061 is again universally conserved and all the three residues are essential for the peptidyl transferase function (Nissen et al., 2000; Polacek and Mankin, 2005).

These interactions have stabilized the N-loop of HflX in a fixed conformation. Intriguingly, all these HflX-interacting PTC residues are also functionally critical, and they mediate the binding of tRNAs, the nascent chain and translation factors to the PTC during the translation cycle. This suggests that the N-loop of HflX could have a profound role in translation regulation.

### The N-loop of HflX clashes with several PTC-targeting antibiotics on the 50S subunit

We noticed that the binding position of the N-loop is also the sites of many PTC-targeting antibiotics. For examples, CHL interacts with C2452 and as well as A2602 of the 23S rRNA and conflicts with the aminoacyl moiety of the A-site; lincomycin (Lin) and its derivative clindamycin (Clin) occupy a similar site involving the HflX-interacting G2505 and its neighboring residues and collide with the A-site tRNA (Dunkle et al., 2010; Lin et al., 2018; Matzov et al., 2017). This prompts us to examine whether the N-loop of HflX directly overlaps with these antibiotics at the PTC. Superimposition of several PTC-targeting antibiotics on the 50S-HflX structure indeed revealed a clash with the N-loop (Fig. 3b-d). R153 of HflX overlaps with the benzene ring of CHL, and G154 clashes with the nitroso-group of CHL (Fig. 3b). R153 also directly conflicts with the heterocycle of Lin and Clin (Fig. 3c-d). In contrast to these A-site binding antibiotics, other antibiotics that are closer to the entry of the peptide tunnel are too far to be sterically affected, such as tetracycline (Tet) and erythromycin (Ert) (Fig. 2e-f).

**Fig. 3.**
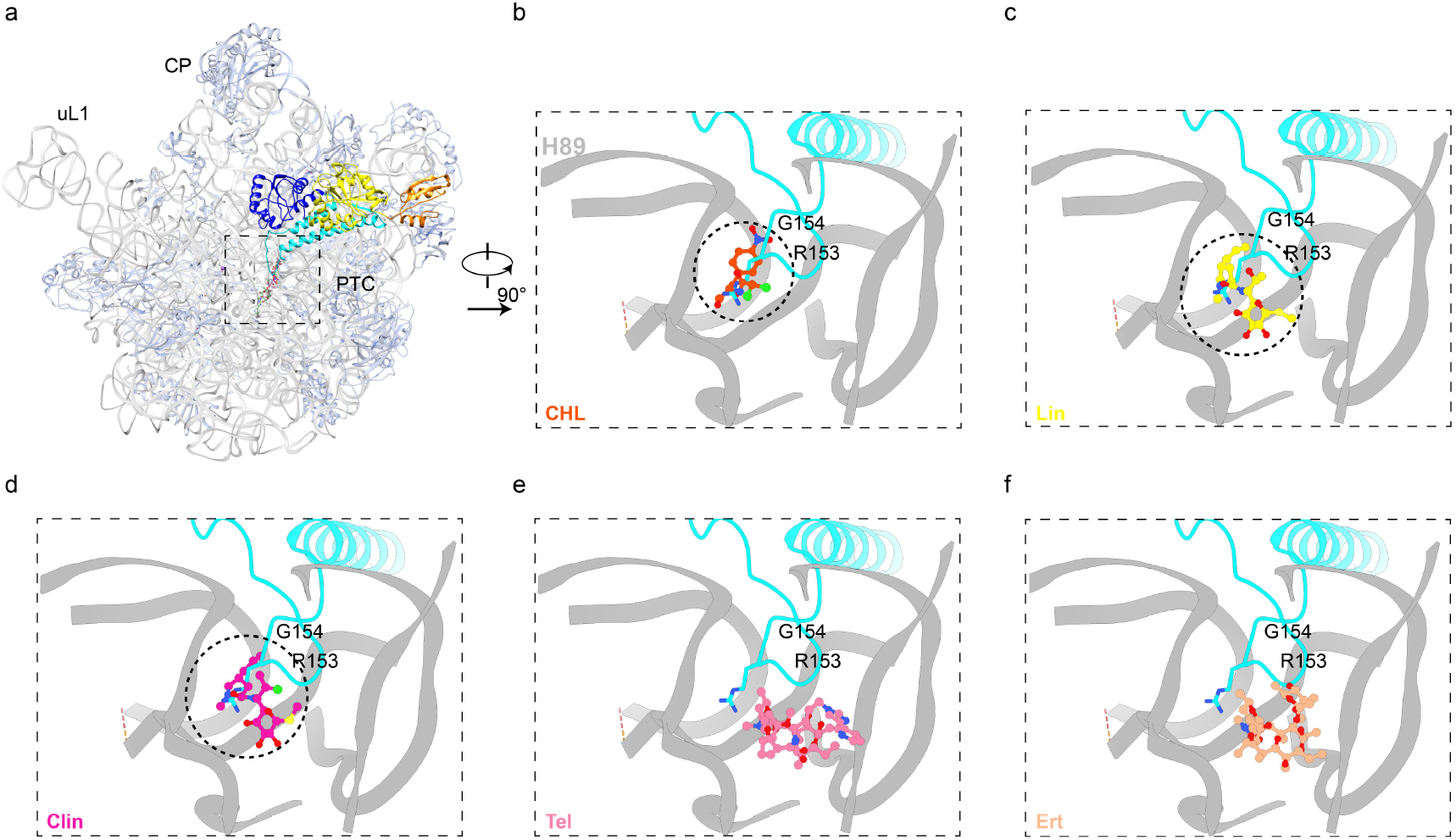
The binding sites of the N-loop of HflX clashes with PTC-targeting antibiotics. **a**, Superimposition of several PTC-targeting antibiotics onto the atomic model of the HflX-50S complex. The chloramphenicol-bound (PDB: 4V7T) (Dunkle et al., 2010), lincomycin-bound (PDB: 5HKV) (Matzov et al., 2017), clindamycin-bound (PDB: 4V7V) (Dunkle et al., 2010), telithromycin-bound (PDB: 4V7S) (Dunkle et al., 2010) and erythromycin-bound (PDB: 4V7U) (Dunkle et al., 2010) 50S or 70S structures were structurally superimposed using the 23S rRNA as the reference for alignment. **b-f**, Comparison of the binding sites of the N-loop and CHL (b), Lin (c), Clin (d), Tel (e) and Ert (f).

These results suggest that the N-loop of HflX upon docking to the PTC could potentially affect the antibiotic binding. HflX from *E. coli* was reported to split the stalled ribosomes induced by heat shock (Zhang et al., 2015). Therefore, it is possible that HflX could also directly contribute to the release of certain PTC antibiotics from the 50S subunit or the 70S ribosome, similar as the tetracycline resistance protein Tet(M) does on Tet arrested ribosomes (Wilson et al., 2020). Since the N-loop is of varying lengths in different bacteria, this possible role of the N-loop might be species-specific and antibiotic specific(Wilson et al., 2020). Indeed, although the N-loop of *E. coli* HflX is not in conflict with Ert (Fig. 3f), the observed Ert resistance in *Listeria monocytogenes* could be resulted from a longer N-loop of its HflX homolog (Duval et al., 2018; Wilson et al., 2020).

### Deletion of *hflX* in *E. coli* results in hypersensitivity to CHL but not to Ert

Following this structural clue, we tested whether *E. coli* HflX could also confer resistance to certain PTC-targeting antibiotics. Firstly, we tested the antibiotics whose binding sites conflict with the N-loop, such as Clin and CHL (Fig. 3a). An antibiotic sensitive assay was carried on with both BW25113 (WT) and JW4131 (Δ*hflX*) strains. Compared to the WT strain, deletion of *hflX* results in a hypersensitive to CHL (Fig. 4a and c), but not to Clin (Fig. 4a). This may due to the fact that Clin is an antibiotic primarily used to inhibit gram-positive bacteria. The sensitivity to CHL was concentration-dependent, and the deletion of *hflX* led to a significant increase of the IC50 from 1.62 μg/mL to 1.1μg/mL (Fig. 4c-d). In contrast, these two strains showed no difference in growth at different concentrations of Ert, even at a very high concentration of the drug (Fig. 4b). The IC50 of Ert to the WT and mutant strains are 43.31 μg/mL and 40.17 μg/mL, respectively, only with a marginal difference (Fig. 3e). This result is in sharp contrast to what has been observed in *Listeria monocytogenes*, indicating that HflX-mediated resistance is indeed species specific. As a control, Tet was assayed for comparison, which targets multiple sites in the 30S subunit. As expected, the two strains displayed little difference under the Tet stress (Supplementary Fig. 3). These results are consistent with the structural comparisons, in which the loop of HflX overlaps the most with the binding of CHL in the PTC (Fig. 3).

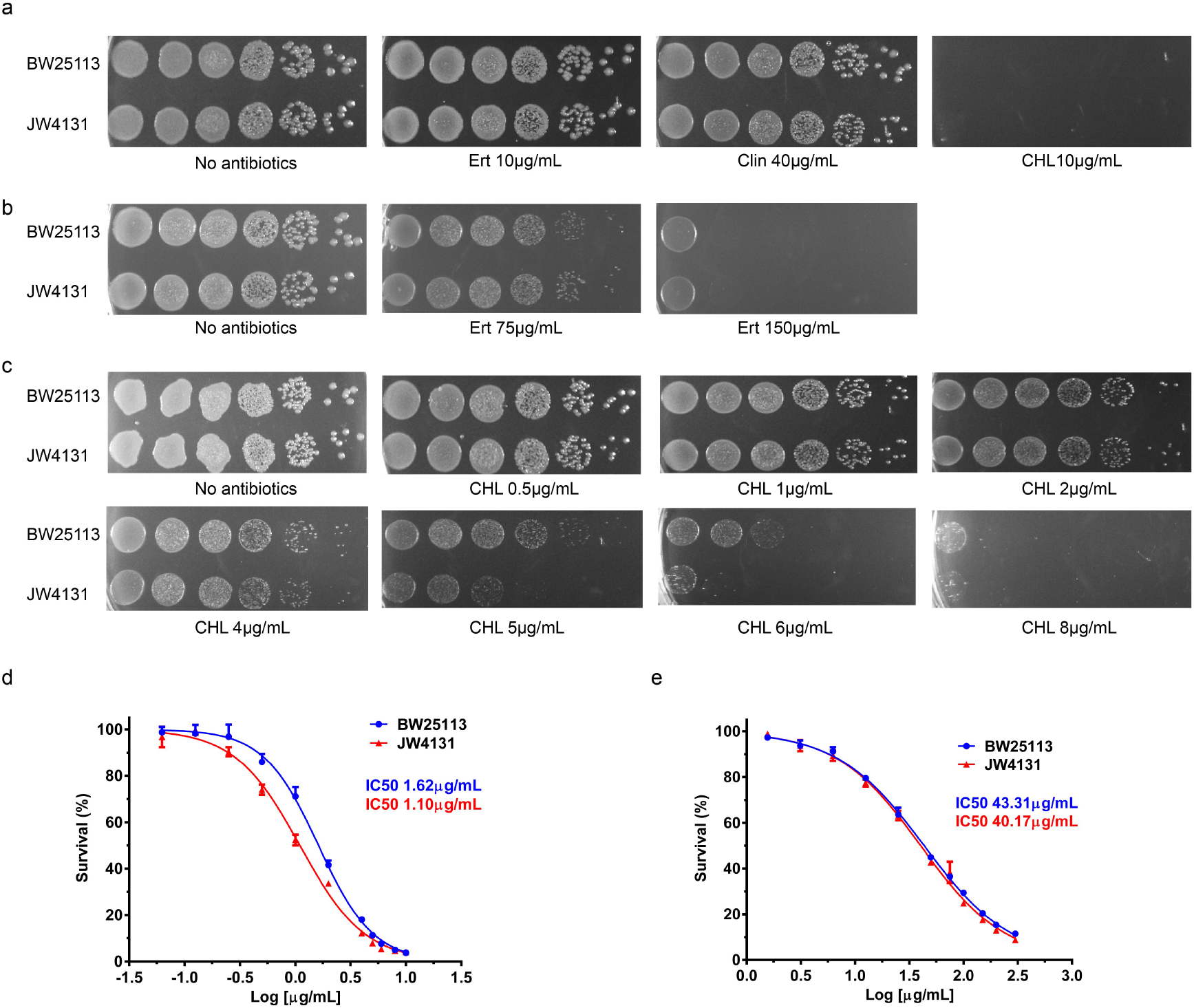
**a**, Ten-fold serial dilutions of WT strain (BW25113) and deletion strain (JW4131) spotted on LB plates containing 20 μg/mL Ert, 40 μg/mL Clin, or 10 μg/mL CHL. **b**, BW25113 and JW4131 cells were spotted on LB plates containing 75 μg/mL or 150 μg/mL Ert. **c**, BW25113 and JW4131 cells were spotted on LB plates containing serial concentrations of CHL from 0.5 to 8 μg/mL. **d**, Growth curves of BW25113 (blue line) and JW4131 (red line) strains with increasing CHL concentration. **e**, Growth curves of BW25113 (blue line) and JW4131 (red line) strains with increasing Ert concentration.

### Stringent control of HflX expression is required for CHL resistance in *E. coli*

To confirm that *hflX* is involved in the CHL resistance, we perform a plasmid-based genetic complementation assay. A pBAD plasmid (pBR322 origin) harboring *hflX* sequence was transformed into the WT and Δ*hflX* strains, and a series of L-arabinose (L-ara) concentrations (0.1% to 1%) were tested. Unexpectedly, under all tested L-ara concentrations, the Δ*hflX* strain grows even worse with the exogenous expression of *hflX* (Supplementary Fig. 4a-c), and the growth inhibition is clearly dose-dependent on L-arabinose. The inhibition on the cell growth is not limited to the Δ*hflX* strain, as the WT strain displays a similar growth defect with the overexpression of *hflX.* We also repeated the experiment by pre-induction of HflX expression for 1 hour before plating, and same results were observed (Supplementary Fig. 4d). It must be noted that without the additional CHL stress, the overexpression of HflX has neglectable effect to the growth of both the WT and Δ*hflX* cells. These data indicate that the expression control of HflX is critical for its function in Chl-induced stress response.

The gene of *hflX* is located in a super-operon (*amiB-mutL-miaA-hfq-hflX-hflK-hflC*) in *E.* coli with a highly complex control of transcription (Tsui et al., 1996; Tsui et al., 1994; Tsui and Winkler, 1994). Three promoters, including both housekeeping σ70 and stress-related σ32, upstream of *hfq* are responsible for the primary transcription of *hflX* in a major unit of *hfq-hflX-hflK-hflC* (Supplementary Fig. 4d). Moreover, *hflX* could be included in much larger transcription units containing several upstream genes and under control of promoters of these genes (Tsui et al., 1996; Tsui et al., 1994; Tsui and Winkler, 1994). On the translation level, the start codon of *hflX* is a less-frequent TTG, and the translational efficiency of ORFs with TTG is seven- to eight-fold lower than that with ATG (Sussman et al., 1996). Therefore, we performed a genetic complementation assay using a genomic fragment containing the upstream regulatory sequence, *hfq* and *hflX* (Supplementary Fig. 4e). This fragment is indeed highly efficient in conferring CHL resistance to the Δ*hflX cells* (Supplementary Fig. 4f).

In summary, our results reveal that as a stress-related factor, the cellular level of HflX is strictly controlled, and excessive HflX is only beneficial under certain stress conditions.

### The N-loop of HflX is essential for the antibiotic resistance

Since the N-loop is in direct conflict with CHL on the 50S subunit, we next assayed whether mutations of certain loop residues could disrupt the HflX-mediated antibiotic resistance in *E. coli.* For this purpose, R153 and G154, which have serious steric clash with CHL (Fig. 3), and their neighboring G151 and P155 were chosen for mutagenesis. Notably, G151 and G154 are also highly conserved across species (Fig. 5a). We also included a loop deletion mutant by removing the nine glycine-rich sequence (K147-G156). We introduced these mutations to the Hfq-HflX plasmid and tested their ability in restoring CHL resistance in Δ*hflX* cells. Among these mutations, the loop deletion is the most effective, followed by G154A mutation, as Δ*hflX* cells harboring these two plasmids are more sensitive to CHL treatment. In contrast, R153A shows no effect, nor do G151A and P155A. The marked effects of G154A and the loop deletion suggest that the N-loop highly likely directly interferes with the binding of CHL to the PTC.

**Fig. 5.**
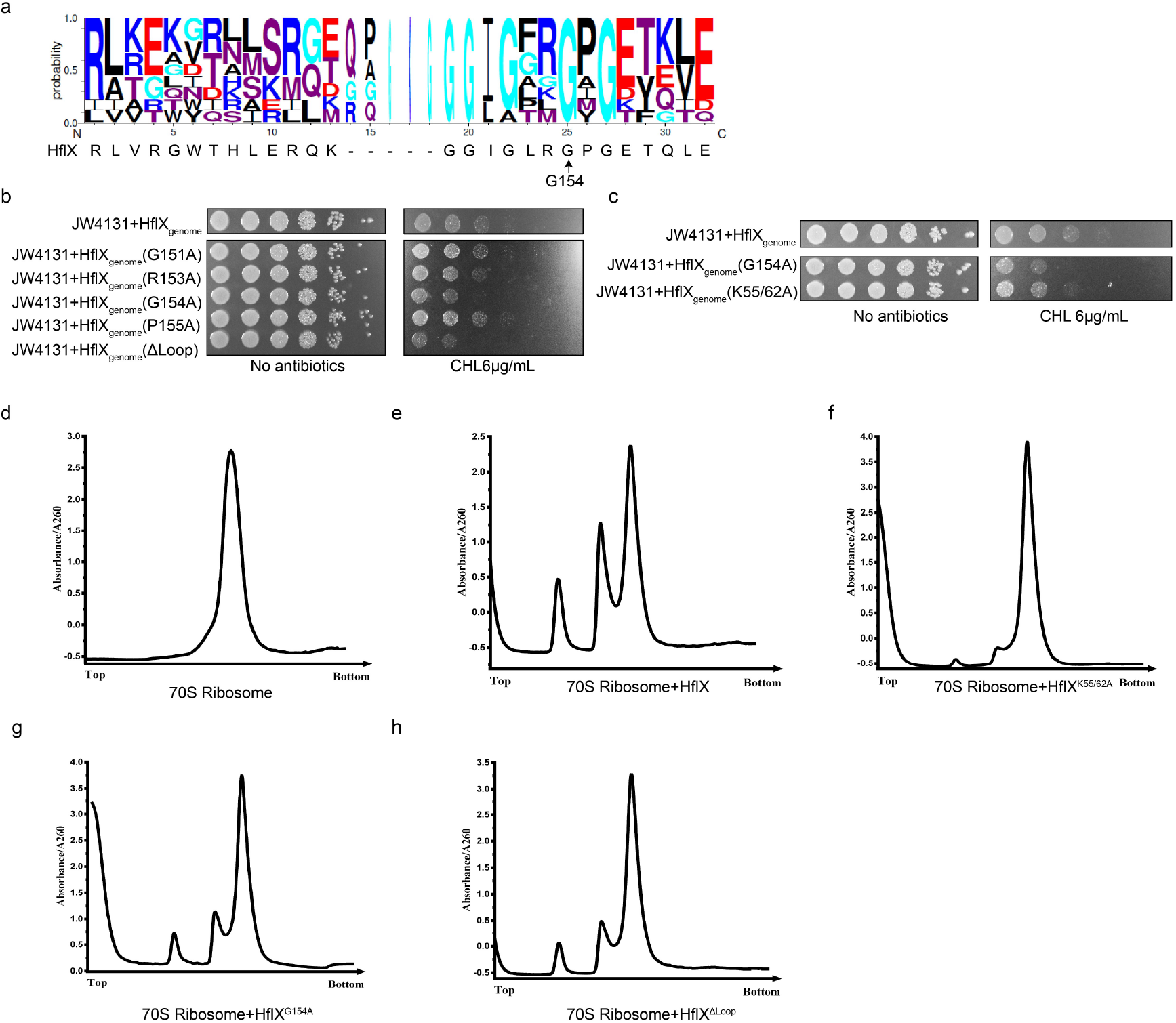
The N-loop of HflX is responsible for HflX-mediated antibiotic resistance. **a**, Sequence conservation of the N-loop of HflX in baceria. G154 is highly conserved among bacteria. **b-c**, Spot assay on the function of HflX mutant variants in CHL resistance. **d-h**, 70S-splitting assay on the ribosome-disassociating activities of HflX variants. 70S ribosomes (0.3 μM) were incubated with various mutants of HflX (6 μM) in the presence of 1 mM GMPPNP at 37°C for 15 min and examined by sucrose density gradient centrifugation.

### Ribosome splitting and CHL displacing are possibly two separate functions of HflX

To explore that the relationship of the CHL resistance and the ribosome-splitting activity of HflX, the recombinantly expressed and purified mutant HflX proteins were subject to an *in vitro* 70S-splitting assay. Consistent with our previous work (Zhang et al., 2015), WT HflX is highly efficient in converting the 70S ribosomes into separate 50S and 30S subunits. In contrast, both the G154A and Δloop mutants exhibit decreased splitting activity. This seems to suggest that the splitting activity and CHL resistance are coupled. Another explanation is that since the N-loop has an extensive interaction with the PTC, it may contribute to the stable binding of HflX to the ribosome to facilitate the splitting. To reconcile this discrepancy, we chose a splitting-defective mutant of HflX (K55A/K62A) (Zhang et al., 2015), in which these two mutations are located in the interface between subdomain 1 and H69/H71, far from the CHL binding site. Consistent with our previous results, the K55A/K62A mutant displays nearly no activity in splitting the 70S ribosome (Fig. 5f). But it is still effective in conferring Chl-resistance to Δ*hflX* cells, significantly stonger than the G154A mutant (Fig. 5c). Thus, the K55A/K62A mutant might have separated the two molecular functions of HflX.

These data suggest that HflX could have two related but mechanistically distinct roles on the ribosome. One is to split the stalled ribosomal complexes accumulated under stress conditions, and the other to dislodge CHL from the 70S ribosome or the 50S subunit. Since CHL is known to induce translational stalling, it is highly possible that the antibiotic displacement could take place during the splitting process.

## Discussion

HflX, a member of the OBG-HflX-like superfamily, is a universally conserved GTPase in three domains of life and plays a vital role in cellular responses to several stresses (Srinivasan et al., 2019; Verstraeten et al., 2011). The gene of *hflX* was found to be located downstream of *hfq* in a super-operon (*amiB-mutL-miaA-hfq-hflX-hflK-hflC*) with a highly complex control of transcription in *E. coli* (Tsui et al., 1996). HflX is under control of heat-induced promoters in the superoperon and has been demonstrated to be a heat-shock protein (Carruthers and Minion, 2009; Chuang and Blattner, 1993; Zhang et al., 2015). HflX was also reported to involve in cellular responses to osmotic and hypoxic stresses (Ngan et al., 2021; Weber et al., 2006), and participates in the regulation of the manganese homeostasis in *E. coli* (Kaur et al., 2014; Sengupta et al., 2018). These data, together with the experiment results that HflX mediates different antibiotic resistance in different bacterial species (Fig. 4) (Duval et al., 2018; Rudra et al., 2020), suggest that HflX could be a general stress response factor, offering protection to bacterial cells from various stresses.

Previously, we have demonstrated that HflX could split stalled ribosomes generated upon heat-shock and by doing so HflX functions to recycle ribosomal subunits to increase the cellular translation capacity to support survival under adverse temperature condition (Zhang et al., 2015). The splitting activity of HflX is evolutionarily conserved in eukaryotes: GTPBP6, a mammalian homolog of HflX in mitochondria, also plays an important role in mito-ribosome recycling (Hillen et al., 2021; Lavdovskaia et al., 2020). Importantly, other types of stresses, such as the antibiotic treatment (Ramu et al., 2009) and osmotic stress (Dai et al., 2018), are known to cause the stalling of elongating ribosomes. Therefore, it is not clear to what extents these various functions of HflX could be attributed to its ribosome-splitting activity.

Based on previous reports on HflX-mediated antibiotic resistance (Duval et al., 2018; Rudra et al., 2020) and the structural observation that the N-loop of HflX locates in the PTC (Zhang et al., 2015), it was proposed that HflX might be directly responsible for antibiotic displacement (Wilson et al., 2020). Those HflX homologs have been named as HflXr (Duval et al., 2018), referring to this resistance phenotype. In the present work, we determined the high-resolution structure of *E. coli* 50S-HflX structure and found that the N-loop of HflX directly clashes with a few PTC-targeting antibiotics. R153 and G154 of HflX spatially overlaps the benzene ring and the nitroso-group of CHL, respectively (Fig. 3). We further demonstrated that the *hflX*-deleted strain of *E. coli* is indeed hypersensitive to CHL, but not to Ert (Fig. 4), which targets the peptide tunnel exit and is relatively away from the N-loop. These results strongly suggest that HflX could potentially directly displace CHL from the 50S subunit. Moreover, combining spot assay and *in vitro* splitting assay, we found that the splitting activity of HflX is generally beneficial for its function in antibiotic resistance. But a splitting-defective mutant (K55A/K62A) shows that the antibiotic resistance phenotype could be separated from its splitting activity, indicating that these two activities of HflX can function independently. Very recently, while our work is under submission, Koller *et al.* determined the cryo-EM structure of *L. monocytogenes* HflXr-50S complex, and also found that the N-loop of HflXr induces position shfits in G2538 (EcG2505) and U2539 (U2506), which is incompatible with the presence of lincomycin (Koller et al., 2022).

Collectively, these recent and our results suggest that the antibiotic resistance function is not limited to certain version of bacterial HflX. The ribosome splitting and antibiotic displacing are two independent and general molecular roles of HflX. The N-loop varies in both the length and amino acid composition in different species (Wilson et al., 2020) (Figure 5), suggesting that divergent evolution of the N-loop might be an adaptive strategy to promote bacterial survival and to combat with competitive antibiotic-producing microbes in different environment.

## Method

### Antibiotic sensitivity assays

WT strains (BW25113) and KO strains (JW4131) of *E. coli* were grown to OD600 of 0.7 at 37 °C, and subjected to 10-fold serial dilutions. 5 μL liquid was spotted on LB plates containing the indicated concentration of antibiotics and incubated overnight at 37 °C.

### Genetic complementation

pBAD vector carried *hflX* gene was transformed to BW25113 or JW4131. The genome fragment containing *hflX* promoter was amplified from BW25113, and cloned into pBAD vector. The site-directed mutation and truncated variants of HflX were introduced by sitespecific mutagenesis. Spotting assays were carried on as described above.

### Protein overexpression and purification

*hflX* was cloned into pET28a vector with a C-terminal His tag. Protein was overexpressed in BL21DE3 cells (*Trans*), and induced with 1mM isopropyl-β-d-thiogalactopyranoside (IPTG) for 18 h at 18 °C. Cells were harvested and resuspended in lysis buffer (50 mM Tris-HCl, pH 8.0, 500 mM NaCl, 5 mM MgCl_2_, and 20 mM imidazole) with 1mM PMSF and 1% Triton X100. Cells were lysed by sonication. The supernatant was incubated with Ni–NTA agarose (GE Healthcare) for 2 h at 4 °C. After washed 3 times with lysis buffer, protein was eluted with elution buffer (50 mM Tris-HCl, pH 8.0, 300 mM NaCl, 5 mM MgCl_2_, and 500 mM imidazole). The protein was loaded onto Superdex75 10/300GL (GE Healthcare), equilibrated with gel filtration buffer (50 mM Tris-HCl, pH 8.0, 300 mM NaCl, 5 mM MgCl_2_ and 5 mM DTT). The site-directed mutation and truncated variants of HflX were introduced by site-specific mutagenesis, and the protein purification method was as same as described above.

### Ribosome purification

*E. coli* 70S and 50S ribosomes were purified as reported previously (Guo et al., 2011; Jelenc, 1980) with some modifications. *E. coli* strain BW25113 cells were harvested after grown to OD600 of 0.7 at 37 °C. For 50S ribosomes purification, cells were lysed in lysis buffer (20 mM Tris-HCl, pH 7.5, 100 mM KCl, 2 mM MgCl_2_ and 2 mM DTT). The supernatant was subjected to sucrose cushion (20 mM Tris-HCl, pH 7.5, 100 mM KCl, 2 mM MgCl_2_, 1 mM DTT and 30% sucrose) in a 70Ti rotor (Beckman Coulter) for 18 h at 30,000 rpm. The pellets were suspended in suspended buffer (20 mM Tris-HCl, pH 7.5, 100 mM KCl, 2 mM MgCl_2_, 1 mM DTT) and incubated at 37 *°C* for 3 h. After centrifuged for 10 min at 15,000 rpm, the supernatant was loaded onto the top of 10%–40% sucrose gradient and centrifuged at 30,000 rpm for 7 h in a SW32 rotor (Beckman Coulter). Fractions containing the 50S ribosomes were collected and buffer changed into lysis buffer. Purification of 70S ribosomes same as described above, especially in buffer containing 10 mM MgCl_2_ always.

### In vitro splitting assay

70S ribosome splitting assay was determined as described previously (Zhang et al., 2015). Purified 70S ribosome (0.3 μM) incubated with HflX (6 μM) in the reaction buffer (20 mM HEPES, pH 7.5, 100 mM NH_4_Cl and 10 mM Mg(OAc)_2_) in the presence of 1mM GMP-PNP. After incubating at 37 °C for 15 min, the reactions were loaded onto a 10%-40% (w/v) sucrose density gradient. The gradients were centrifuged at 39,000 rpm for 3.5 h at 4°C using a Beckman SW41 rotor (Beckman Coulter) followed by a UV monitor and a density gradient fractionator (Teledyne Isco). Point mutants or truncated versions were performed as mentioned above.

### Cryo-EM sample preparation and data collection

50S ribosomes were incubated with purified HflX in reaction buffer in the presence of 1 mM GMP-PNP for 15 min at 37 °C. Before preparing grids for cryo-EM, sample was centrifuged at 13,000 g for 10 min to remove protein aggregates. 4 μL of reaction complex was loaded on glow-discharged Quantifiol R1.2/1.3 copper grids (Cu 400) covered with a 2-nm thickness carbon made in-house, and the cryo-grids were prepared using a FEI Vitrobot (Mark IV) at 6°C and 100% humidity. Cryo-EM samples were checked with 200 kV FEI Talos Arctica (FEI Ceta camera) before data collection.

Micrographs were recorded with 300 kV FEI Titan Krios TEM (Gatan K2 summit camera) with GIF Quantum energy filter (Gatan). All the data was collected at the magnification of 130,000× and at pixel sizes of 1.052 Å. 32 frames with exposure time of 8 s were collected with defocus ranged from −0.7 to −1.2 μm under low-dose model. A total of 2,304 micrographs were collected with serial EM (Table S1).

### Image processing

Cryo-EM data were subjected to beam-induced motion correction using MotionCor2 (Zheng et al., 2017), and contrast transfer function (CTF) parameters for each micrograph were determined by Gctf (Zhang, 2016). RELION (versions 3.1) was used to perform data processing (Zivanov et al., 2020). 297,527 particles were auto-picked from 2,288 micrographs using RELION 3.1, and subjected to 2D classification. 283,432 particles, selected from 25 classes, were used for further processing. 145,641 particles were applied for 3D refinement, and then re-extracted at pixel sizes of 1.052 Å, after the first round of 3D classification. 93,913 particles were selected and applied for the second round of 3D refinement, after the second round of 3D classification. The final round of refinement was performed with a soft-edged mask applied, resulting in a 3.12 Å map. A 2.84 Å structure was determined after postprocession option. After correction for the modulation transfer function of the K2 detector, CTF refinement and Bayesian polishing, the overall resolution of the final density map was improved to 2.52 Å when particles were refined with a soft-edged mask. The post-processing was done either by RELION3.1 (Zivanov et al., 2020) or DeepEMhancer (Sanchez-Garcia et al., 2021).

### Model building and refinement

Model building was based on the structure of ribosome (Stojkovic et al., 2020) (PDB code: 6PJ6). The existing ribosome model and HflX (Zhang et al., 2015) (PDB code: 5ADY) were first docked into the cryo-EM density map using UCSF Chimera (Pettersen et al., 2004), and then rebuilt manually in Coot (Emsley and Cowtan, 2004), and refined (real-space) using Phenix (Afonine et al., 2018). Figure preparation and structure analysis were performed with and UCSF ChimeraX (Pettersen et al., 2021) and Chimera (Pettersen et al., 2004).

## Data availability

The cryo-EM density maps have been deposited in the Electron Microscopy Data Bank with the accession codes EMDB-33904 [https://www.ebi.ac.uk/emdb/entry/EMD-33904]. The atomic model has been deposited in Protein Data Bank with accession code PDB-7YLA [http://doi.org/10.2210/pdb7ykl/pdb].

## Acknowledgments

We thank the Core Facilities at the School of Life Sciences, Peking University for help with negative staining EM; the Electron Microscopy Laboratory and Cryo-EM Platform for help with data collection; the High-performance Computing Platform for help with computation; the National Centre for Protein Sciences at Peking University for assistance with mass spectrum. The work was supported by the National Science Foundation of China (31725007 to N.G., 31922036 to N.L.), the National Key Research and Development Program of China (2019YFA0508904 to N.G.), and the Qidong-SLS Innovation Fund to N.G.

## Author contributions

D.W. and Y.H. prepared the protein samples. D.W. collected the cryo-EM data, performed EM analysis. D.W. carried out functional experiments under the supervision of N.G.. N.G. and D.W. performed model building and wrote the manuscript.

## Competing interests

Authors declare no competing interests.

**Correspondence and requests for materials** should be addressed to N.G.

---

**Supplementary Fig. 1.**
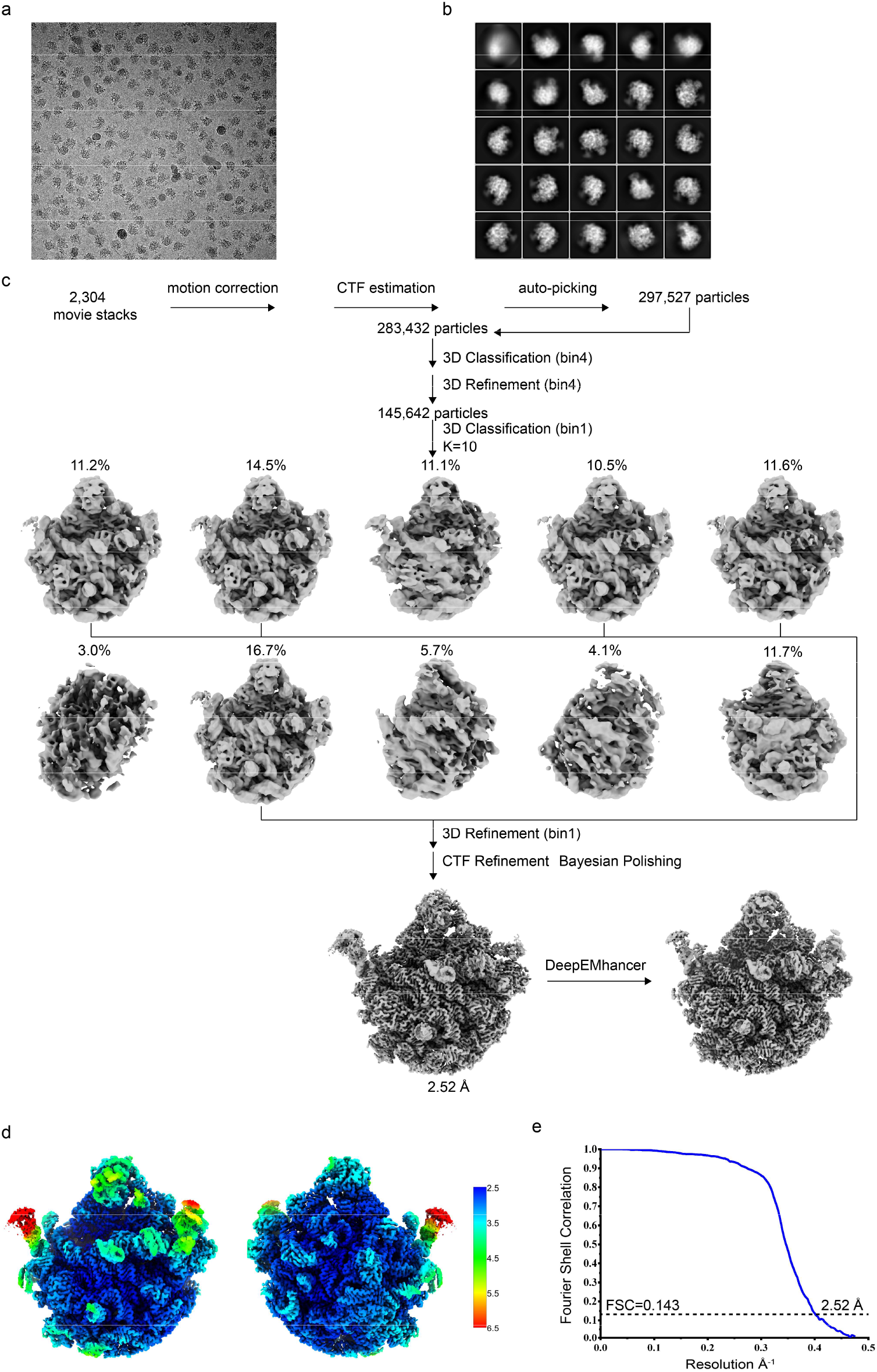
Image processing workflow of the 50S-HflX dataset. **a**, A representative raw cryo-EM micrograph. **b**, Two-dimensional class averages of the HflX-50S particles. **c**, Flowchart of the image processing. **d**, Local resolution estimation of the final cryo-EM map. **e**, Fourier Shell Correlation (FSC) curve for the final cryo-EM map using the gold-standard FSC 0.143 criteria.

**Supplementary Fig. 2.**
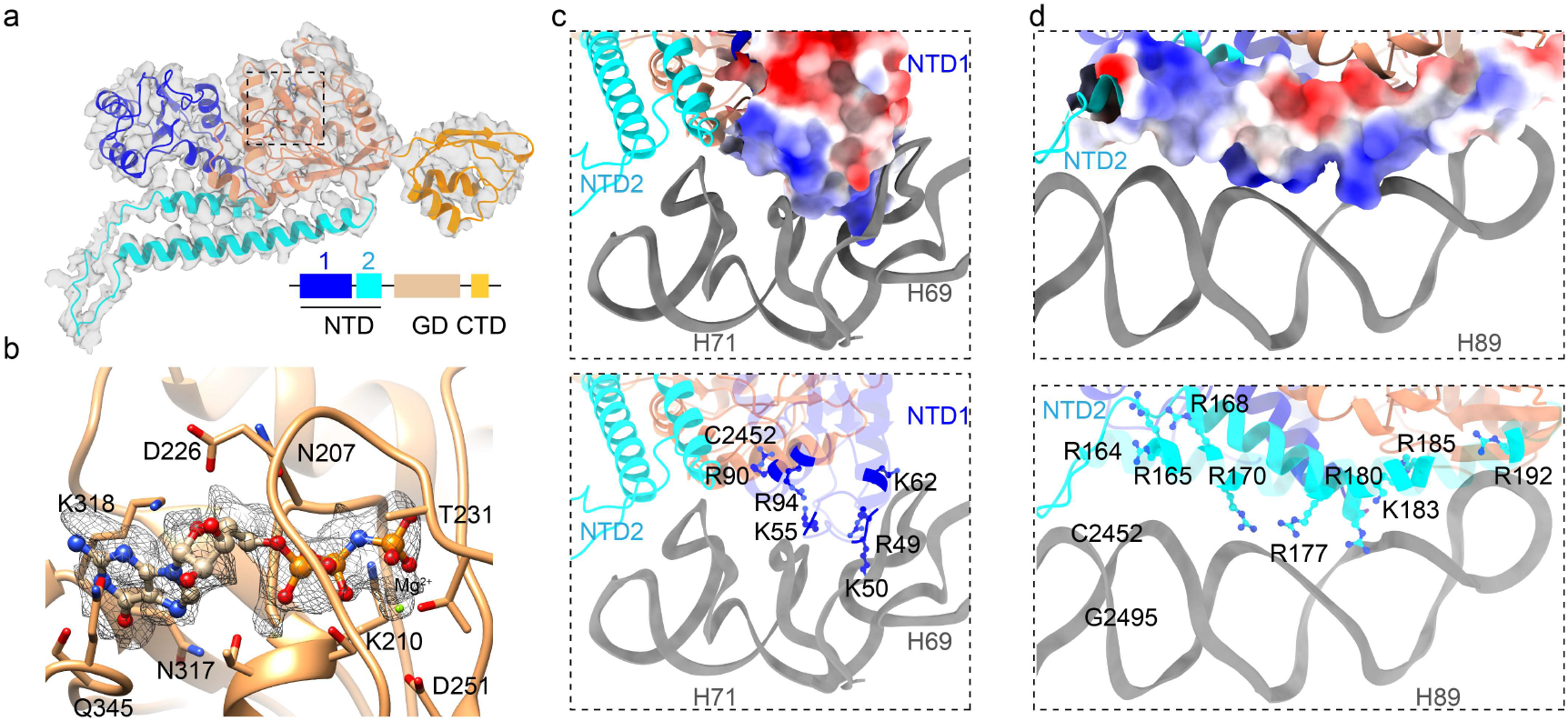
Specific interactions between the 50S subunit and HflX. **a**, The atomic model and segmented density map of HflX. NTD, N-terminal domain (colored in blue and cyan); GD, GTPase domain (colored in yellow); CTD, C-terminal domain (colored in orange). **b**, Zoom-in views of the densities of GMP-PNP. **c**, The interactions between H69-H71 and the NTD of HflX. A positively charged surface patch from the subdomain 1 interacts extensively with the H69 and H71. **d**, The interactions between H89 and the NTD of HflX. The basic residues, including R164, R165, R170, R177, K183, R180 and R192 in the subdomain 2, lie in the interface and interacts extensively with H89.

**Supplementary Fig. 3.**
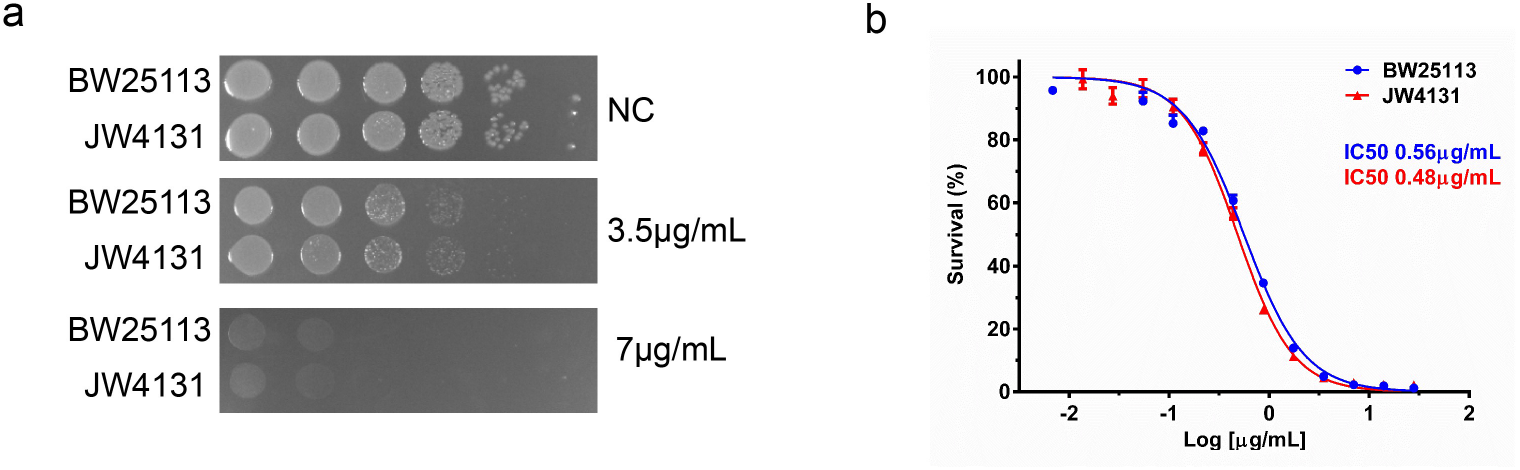
Deletion of HflX is not sensitively to tetracycline. **a**, Ten-fold serial dilutions of WT strain (BW25113) and deletion strain (JW4131) spotted on LB plates containing different concentrations of tetracycline. **b**, Growth curves of BW25113 (blue line) and JW4131 (red line) strains with increasing Tet concentration.

**Supplementary Fig. 4.**
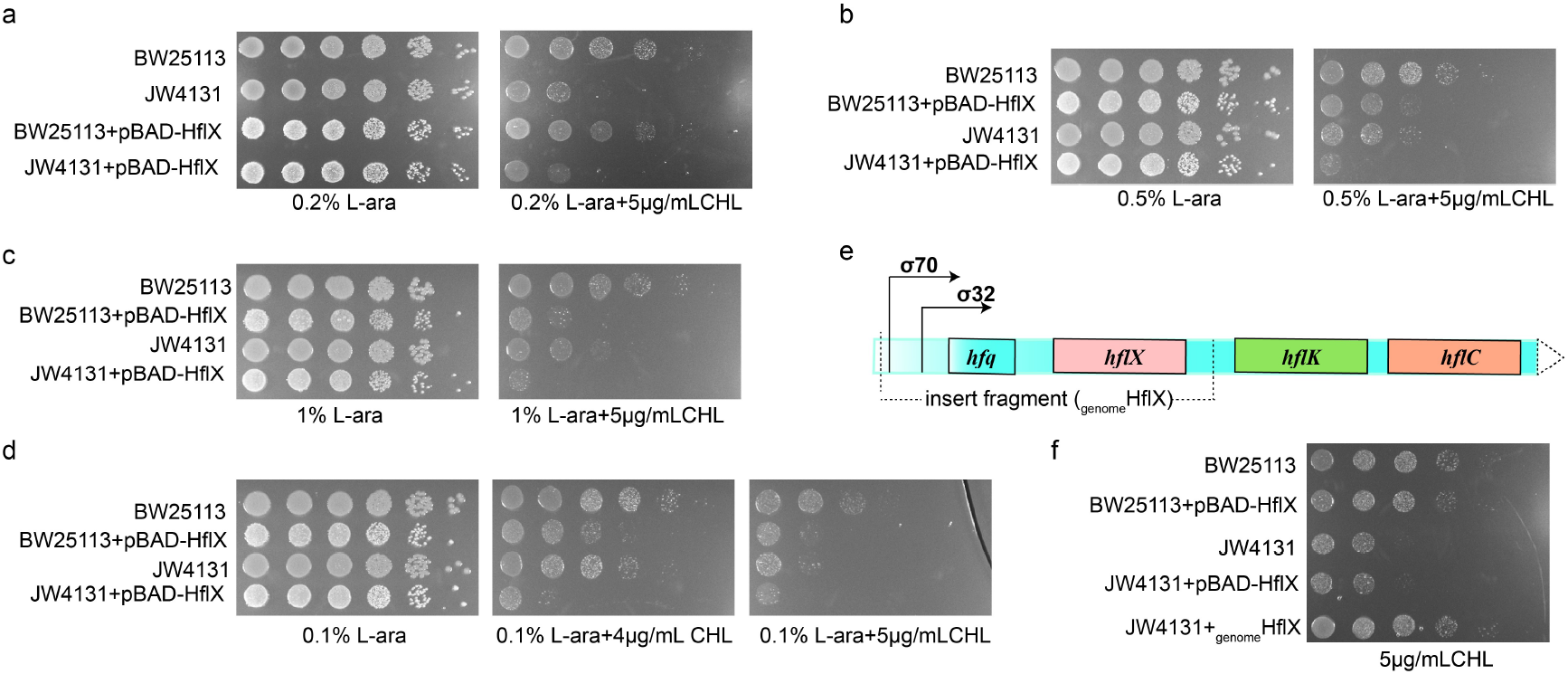
Excessive HflX is not beneficial to bacterial growth under stress conditions. **a-d**, Spot assay of *E. coli* strains (BW25113, BW25113+pBAD-HflX, JW4131, JW4131+ pBAD-HflX) with or without 5 μg/mL CHL in different L-ara concentrations. **e**, Schematic representation of the *hfq-hflX-hflK-hflC* operon under the control of σ^**32**^ and σ^**70**^. The insert fragment indicates the sequence used for genetic complementation experiments in Figure 5. **f**, BW25113, BW25113+pBAD-HflX, JW4131, JW4131+pBAD-HflX and JW4131+ HflX_genome_ stains were spotted on LB plate containing 5 μg/mL Chl.

**Table S1.** Cryo-EM data collection, refinement and validation statistics.

|  |  |
| --- | --- |
|  | 50S-HflX<br>(EMDB-33904)<br>(PDB 7YLA) |
| **Data collection and processing** |  |
| Magnification | 36,000× |
| Voltage (kV) | 300 |
| Electron exposure (e-/Å<sup>2</sup>) | 64 |
| Defocus range (μm) | -0.7 to -1.2 |
| Pixel size (Å) | 1.052 |
| Symmetry imposed | C1 |
| Final particle images (no.) | 93,913 |
| Map resolution (Å) | 2.52 |
| FSC threshold | 0.143 |
| **Refinement** |  |
| Initial model used (PDB code) | 6PJ6 |
| Model resolution (Å) | 2.6 |
| FSC threshold | 0.143 |
| Map sharpening *B* factor (Å<sup>2</sup>) | -79 |
| Model composition |  |
| Non-hydrogen atoms | 125,963 (32,077) |
| Protein residues | Protein: 3,775<br>Nucleotide: 3,001 |
| Ligands | Mg<sup>2+</sup>, GNP, Zn<sup>2+</sup> |
| *B* factors (Å<sup>2</sup>) |  |
| Protein | 53.35 |
| Nucleotide | 54.61 |
| Ligand | 76.61 |
| R.m.s. deviations |  |
| Bond lengths (Å) | 0.008 |
| Bond angles (°) | 0.727 |
| Validation |  |
| MolProbity score | 1.77 |
| Clashscore | 7.96 |
| Rotamers outliers (%) | 0.07 |
| Ramachandran plot |  |
| Favored (%) | 95.20% |
| Allowed (%) | 4.72% |

